# The SARS-CoV-2 Spike protein disrupts human cardiac pericytes function through CD147-receptor-mediated signalling: a potential non-infective mechanism of COVID-19 microvascular disease

**DOI:** 10.1101/2020.12.21.423721

**Authors:** Elisa Avolio, Michele Carrabba, Rachel Milligan, Maia Kavanagh Williamson, Antonio P Beltrami, Kapil Gupta, Karen T Elvers, Monica Gamez, Rebecca Foster, Kathleen Gillespie, Fergus Hamilton, David Arnold, Imre Berger, Massimo Caputo, Andrew D Davidson, Darryl Hill, Paolo Madeddu

## Abstract

Severe coronavirus disease 2019 (COVID-19) manifests as a life-threatening microvascular syndrome. The severe acute respiratory syndrome coronavirus 2 (SARS-CoV-2) uses the Spike (S) protein to engage with its receptors and infect host cells. To date, it is still not known whether heart vascular pericytes (PCs) are infected by SARS-CoV-2, and if the S protein alone provokes PC dysfunction. Here, we aimed to investigate the effects of the S protein on primary human cardiac PC signalling and function. Results show, for the first time, that cardiac PCs are not permissive to SARS-CoV-2 infection *in vitro*, whilst a recombinant S protein alone elicits functional alterations in PCs. This was documented as: (1) increased migration, (2) reduced ability to support endothelial cell (EC) network formation on Matrigel, (3) secretion of pro-inflammatory molecules typically involved in the *cytokine storm*, and (4) production of pro-apoptotic factors responsible for EC death. Next, adopting a blocking strategy against the S protein receptors angiotensin-converting enzyme 2 (ACE2) and CD147, we discovered that the S protein stimulates the phosphorylation/activation of the extracellular signal-regulated kinase 1/2 (ERK1/2) through the CD147 receptor, but not ACE2, in PCs. The neutralisation of CD147, either using a blocking antibody or mRNA silencing, reduced ERK1/2 activation and rescued PC function in the presence of the S protein. In conclusion, our findings suggest that circulating S protein prompts vascular PC dysfunction, potentially contributing to establishing microvascular injury in organs distant from the site of infection. This mechanism may have clinical and therapeutic implications.

**Clinical perspective:**

- Severe COVID-19 manifests as a microvascular syndrome, but whether SARS-CoV-2 infects and damages heart vascular pericytes (PCs) remains unknown.
- We provide evidence that cardiac PCs are not infected by SARS-CoV-2. Importantly, we show that the recombinant S protein alone elicits cellular signalling through the CD147 receptor in cardiac PCs, thereby inducing cell dysfunction and microvascular disruption *in vitro*.
- This study suggests that soluble S protein can potentially propagate damage to organs distant from sites of infection, promoting microvascular injury. Blocking the CD147 receptor in patients may help protect the vasculature not only from infection, but also from the collateral damage caused by the S protein.

## INTRODUCTION

Severe coronavirus disease 2019 (COVID-19) is considered a microvascular disorder in which the infectious agent harms vascular cells, causing inflammation and thrombosis.(1–3) Thromboembolic events resulting in stroke or myocardial infarction occur in up to 4% of patients with COVID-19 hospitalised in intensive care units. Moreover, people with pre-existing cardiovascular disease are more likely to die of COVID-19.(4)

Severe acute respiratory syndrome coronavirus 2 (SARS-CoV-2) uses the homotrimeric spike (S) glycoprotein embedded in the virus to bind to cognate receptors on human cells. Such binding triggers a cascade of events that leads to fusion of the viral and cellular membranes to facilitate virus entry,(5) and subsequent manipulation of the host Raf/MEK/ERK signalling pathway to regulate viral replication and gene transcription in host cells.(6) The receptor-binding domain (RBD) contained in the S1 subunit of the viral S protein recognises and binds to the host receptor angiotensin-converting enzyme 2 (ACE2), while the S2 subunit mediates viral - cell membrane fusion by forming a six-helical bundle *via* the two-heptad repeat domain.(7, 8) Interestingly, a recombinant S1 subunit (Val16 – Gln690), induced MEK phosphorylation in pulmonary vascular cells which did not occur when a truncated S1 subunit (Arg319 – Phe541) containing the ACE2 RBD was used.(9) This finding suggests that the viral protein can induce intracellular signalling through alternative receptors, independently of ACE2. Accordingly, another report indicated that the S2 subunit was responsible for disturbing the barrier function of brain endothelial cells (ECs).(10) Moreover, a preclinical study showed that neither soluble ACE2 nor ACE2 substrates prevented the cellular uptake of the S1 subunit in selected non-pulmonary tissues, implying the presence of alternative receptors to ACE2.(11)

CD147, also known as Basigin or extracellular matrix metalloproteinase inducer (EMMPRIN), has recently emerged as a novel receptor for SARS-CoV-2.(12) It was previously identified as an entry receptor for measles virus on epithelial cells.(13) This transmembrane protein is expressed by ECs, signals through ERK1/2, is upregulated during inflammation and atherothrombosis, and may contribute to plaque instability by inducing metalloproteinase expression.(14) Therefore, CD147 represents a potential mediator of the cardiovascular damage caused by SARS-CoV-2. It remains, however, unknown whether SARS-CoV-2 infects and harms vascular cells of the heart through a mechanism involving ACE2 or Basigin/CD147, or both. Whilst the expression of ACE2 in ECs remains uncertain, (15–18) the recognition of elevated ACE2 mRNA transcript levels suggested a predominant functional role for ACE2 in coronary mural cells.(19, 20)

Pericytes (PCs) are pleiotropic cells that wrap ECs. In the heart, they are abundantly associated with the coronary microvasculature.(21) They support the integrity of coronary artery ECs (CAECs),(22, 23) participate in vascular remodelling(24) and cardiac repair,(25) and modulate inflammatory responses.(26) Dysfunctional cardiac PCs were found in patients with severe myocardial disease.(27) Dysfunctional PCs participate in adverse vascular phenomena; for instance, after a heart attack, persistently contracted PCs block the coronary microvascular circulation, thereby causing blood to clot.(28) Interestingly, a reduction in the vascular coverage by PCs was documented in the heart and lungs of human patients with COVID-19, in the absence of capillary rarefaction,(29, 30) suggesting that SARS-CoV-2 may affect the microvasculature by specifically targeting PCs. Therefore, it is crucial to understand the role of these perivascular cells in COVID-19.

The aims of this study were to: (1) explore if human cardiac PCs express ACE2 or CD147, or both; (2) verify if PCs are permissive to SARS-CoV-2 infection *in vitro*; (3) investigate whether a recombinant SARS-CoV-2 S protein alone, outside the context of infectious virus, can trigger molecular, functional, and pro-inflammatory alterations in PCs; (4) To adopt blocking antibodies or mRNA silencing strategies to understand which viral receptor is responsible for the harmful S protein effects in PCs.

## METHODS

Unless otherwise stated, all chemicals were purchased from Sigma-Aldrich, UK.

### Ethics

This study complies with the ethical guidelines of the Declaration of Helsinki.

#### Histology on human hearts

myocardial samples collection was covered by the Independent Ethics Committee of the University Hospital of Udine (October 22nd, 2013, ref. 58635). Patients (N=3) were enlisted for cardiac transplant or device implantation due to end stage heart failure. All patients signed informed consent. Patients were recruited before the COVID-19 pandemic.

#### Extraction of primary cardiac PCs

human myocardial samples were discarded material from surgical repair of congenital heart defects (ethical approval number 15/LO/1064 from the North Somerset and South Bristol Research Ethics Committee). Adult patients and paediatric patients’ custodians gave informed written consent. Donors and samples characteristics are described in **Table 1**. Patients were recruited before the COVID-19 pandemic.

**Table 1.** Cardiac pericytes donors.

| Sample type | Congenital heart defect / intervention needed | Patient age |
| --- | --- | --- |
| RV | Pulmonary atresia | 11 days |
| RA | Total anomalous pulmonary vein connection | 18 days |
| RV | Ventricular septal defect | 6 months |
| RV | Tetralogy of Fallot | 6 months |
| RV | Atrial-Ventricular canal closure | 3 years |
| RV | Pulmonary valve repair | 14 years |
| RV | Pulmonary valve repair | 15 years |
| RA | Atrial septal defect | 17 years |
| RV | Tricuspid valve replacement + Pulmonary valve repair | 23 years |
| RA | Atrial septal defect | 54 years |

RA: right atrium
RV: right ventricle

#### Serological studies

serum samples from COVID-19 patients (N=64; 32M/32F; age range 20-93 years) were collected as part of the DISCOVER study from patients admitted to North Bristol NHS Trust (Ethics approval via South Yorks REC: 20/YH/0121, CRN approval no: 45469). Blood was withdrawn between 0 and 34 days from the onset of COVID-19 symptoms. Pre-pandemic control sera (N=14; 3M/11F; age range 30-69 years) were randomly selected from 526 anonymised blood donor samples (age range 18-69 years; 277M/249F). Ethical approval number 19/WA/0295 was granted by Wales 6 REC. All donors and patients gave informed written consent.

### Immunofluorescence analysis of cardiac PCs *in situ*

Human myocardial samples were either fixed in formalin and paraffin embedded, or frozen using OCT compound. 5-μm thick sections were cut for identification of cardiac PCs *in situ*. Paraffin sections required heat-induced antigen retrieval, performed using citrate buffer 0.01M pH=6, for 40 min at 98 °C. Tissue sections were blocked with 10% v/v normal donkey serum and incubated with primary antibodies for 16h at 4°C. Antibodies were: anti-platelet derived growth factor receptor beta (PDGFRβ - R&D AF385, 1:50), anti-smooth muscle alpha-actin (α-SMA - Dako GA611, 1:100), anti-ACE2 (Merck SAB3500346, 1:40), anti-CD147 (BioLegend #306221, 1:100), anti-von Willebrand Factor (vWF - Merck F3520, 1:200). Secondary antibodies (Alexa 488-, Alexa 568-, Alexa 647-conjugated) were all purchased from ThermoFisher Scientific and used at a dilution of 1:200, for 1h at 20°C in the dark. Slides were mounted using ProLong™ Gold Antifade Mountant with 4’,6-diamidino-2-phenylindole (DAPI) (ThermoFisher Scientific). Imaging was performed using a Leica TCS SP8 confocal microscope.

### Primary cultures of cardiac PCs and ECs

Immunosorted PCs were expanded in a dedicated medium (ECGM2, C-22111, PromoCell) supplemented with human recombinant growth factors and 2% v/v foetal bovine serum (FBS), as previously described.(22, 31) Human coronary artery ECs (CAECs) were purchased from PromoCell and expanded in the same medium used for PCs. All cells used in this study tested free of mycoplasma contamination. Cells were used between passage 4 and 7.

### Cell line culture

The human gut epithelial cell line, Caco2, expressing hACE2 (Caco-2-ACE2) was a kind gift from Dr Yohei Yamauchi, University of Bristol. The African green monkey kidney cell line VeroE6 engineered to overexpress the human ACE2 and TMPRSS2 (VeroE6/ACE2/TMPRSS2)(32) was a kind gift from Dr Suzannah Rihn, MRC-University of Glasgow Centre for Virus Research. All cells were cultured in Dulbecco’s modified Eagle’s medium plus GlutaMAX (DMEM, Gibco, ThermoFisher) supplemented with 10% v/v FBS, 1% v/v sodium pyruvate, and 0.1 mM non-essential amino acids.

The human lung epithelial cell line Calu3 (ATCC HTB-55) was cultured in Eagle’s minimum essential medium plus GlutaMAX (MEM, Gibco, ThermoFisher) with 10% v/v FBS, 0.1 mM non-essential amino acids, and 1% v/v sodium pyruvate.

### Immunocytochemistry (ICC) analyses

Cells were rinsed with PBS and fixed with 4% w/v paraformaldehyde (PFA) in PBS for 15 min at 20°C. After washing with PBS, the cells were permeabilised with 0.1% v/v Triton-X100 in PBS for 5 min at 20°C, when required. Cells were blocked with 10% v/v FBS and incubated with the following antibodies for 16 hrs at 4°C: anti-ACE2 (R&D AF933, dilution 1:50); anti-CD147 (BioLegend #306221, 1:100); anti-Transmembrane Serine Protease 2 (TMPRSS2 - Proteintech 14437-1-AP, 1:100); anti-Neural/Glial antigen 2 (NG2 - Millipore AB5320, 1:100); anti-PDGFRβ (R&D AF385, 1:100); anti-PDGFRα (Santa Cruz sc-398206, 1:100). Secondary antibodies conjugated with either Alexa 488 or Alexa 568 or Alexa 647 were purchased from ThermoFisher Scientific and used at a dilution of 1:200, for 1 h at 20°C, in the dark. Nuclei were counterstained using DAPI. Images were snapped and processed using a Zeiss AxioObserver Z1 Microscope equipped with a 20x objective.

### Production and purification of recombinant SARS-CoV-2 S protein

SARS-CoV-2 S protein was expressed in insect cells and purified as described previously.(33, 34) Briefly, the S construct encoded amino acids 1 to 1213 (extracellular domain) fused with a thrombin cleavage site, followed by a T4-foldon trimerisation domain and a hexahistidine affinity purification tag at the C-terminus. The polybasic furin cleavage site was mutated (RRAR to A) to increase the stability of the protein for *in vitro* studies.(33, 34) S protein was expressed in Hi5 cells using the MultiBac system.(35) Secreted S protein was harvested 3 days after infection by centrifuging the cell culture at 1,000*g* for 10 min followed by another centrifugation of supernatant at 5,000*g* for 30 min. S protein-containing medium was incubated with HisPur Ni-NTA Superflow Agarose (Thermo Fisher Scientific) for 1h at 4°C. Resin bound with S protein was separated from unbound proteins and medium using a gravity flow column, followed by 30 column volume wash with wash buffer (65 mM NaH_2_PO_4_, 300 mM NaCl, 20 mM imidazole, pH 7.5). Finally, the protein was eluted with a step gradient of elution buffer (65 mM NaH_2_PO_4_, 300mM NaCl, 235mM imidazole, pH 7.5). Eluted fractions were analysed by reducing SDS-PAGE. Fractions containing the S protein were pooled and concentrated using 50 kDa MWCO Amicon centrifugal filter units (EMD Millipore). During concentration, proteins were buffer-exchanged in phosphate-buffered saline (PBS) pH 7.5. Concentrated protein was aliquoted, flash frozen in liquid nitrogen, and stored at −80°C until use.

### Measurement of S protein in patients’ sera

The presence of S protein in COVID-19 patients’ serum was evaluated using the COVID-19 Spike Protein ELISA Kit from Abcam (ab274342), according to manufacturer’s instructions. Pre-pandemic sera were employed as controls. All test sera were diluted 1:2. The S protein concentration was expressed as ng per mL serum.

### Western blotting on total cell lysates

Whole-cell protein lysates were prepared using RIPA buffer supplemented with 1:50 proteases inhibitors cocktail and 1:100 phosphatases inhibitors. Protein extracts were centrifuged 15 min at 10,000 *g*, 4°C. After the assessment of protein concentration (BCA Protein Assay Kit, ThermoFisher Scientific), the supernatants were kept at −80°C. Protein samples (10-15 μg) were prepared in Laemmli loading buffer, incubated for 8 min at 98°C, resolved on 10% SDS-PAGE, and transferred onto 0.2 μm PVDF membrane (Bio-Rad). Membranes were blocked using 5% w/v non-fat dried milk (Bio-Rad) in Tris-buffered saline (TBS, BioRad) supplemented with 0.05% v/v Tween-20 for 2 h at 20°C. Primary antibodies (ACE2, dilution 1:100; TMPRSS2, 1:500; CD147, 1:500; 6x-HIS-tag (Invitrogen MA1-21315), 1:1000; P-ERK1/2 Thr202/Tyr204 (Cell Signaling Technology #4370), 1:2000; Total ERK1/2 (Cell Signaling Technology #4395), 1:1000) were incubated for 16 hrs at 4°C. GAPDH was used as a loading control (Cell Signaling Technology #97166, 1:1000). Anti-mouse or anti-rabbit IgG HRP (1:5000, both from GE Healthcare) or anti-goat IgG HRP (R&D HAF017, 1:5000) were employed as secondary antibodies. Membrane development was performed by an enhanced chemiluminescence-based detection method (ECL™ Prime Western Blotting Detection Reagent, GE Healthcare) and observed using a ChemiDoc MP Imaging System (Bio-Rad). Western blot data were analysed using the BioRad Image Lab and the ImageJ software.

For detection of the S protein binding to PCs, 1 μg/mL (5.8 nM) S protein or PBS vehicle were incubated with PCs for 1 h at 37°C, and whole cell protein lysates collected in RIPA buffer as described.

### Infections of primary cardiac PCs and Caco-2 cells with SARS-CoV-2

Stocks of SARS-CoV-2 viral isolates, SARS-CoV-2/human/Liverpool/REMRQ0001/2020 (REMRQ0001, wild strain) or hCoV-19/England/204690005/2020 (lineage B.1.1.7 - alpha variant; GISAID ID: EPI_ISL_693401; kindly provided by Professor Wendy Barclay, Imperial College, London and Professor Maria Zambon, Public Health England) were produced by inoculation of VeroE6/TMPRSS2 cells and titred as previously described.(33, 36) Caco-2-ACE2 cells and cardiac PCs were plated in μClear 96-well microplates (Greiner Bio-one) and the next day infected with either REMRQ0001 or B.1.1.7 at multiplicity of infection (MOI) of 10, using respective culture media, and incubated at 37°C. Uninfected controls which received media only were also included.

After 24 hrs, cells were fixed with 4% w/v PFA for 60 min before being permeabilised with 0.1% v/v Triton-X100 and blocked with 1% v/v bovine serum albumin (BSA). Cells were stained with DAPI and antibodies against double-stranded RNA (dsRNA, J2 10010200, Scicons) and SARS-CoV-2 nucleocapsid (N) protein (200-401-A50, Rockland) followed by appropriate secondary antibodies conjugated with Alexa Fluor dyes 586 and 647. An ImageXpress Pico Automated Cell Imaging System (Molecular Devices) was used to capture immunofluorescence using a 10X objective. All work with infectious SARS-CoV-2 virus was conducted in a Class III microbiological safety cabinet in a containment level 3 facility at the University of Bristol.

### Assessment of cell viability

Viability of cardiac PCs and CAECs exposed to 1 μg/mL (5.8 nM) S protein or PBS vehicle was evaluated using the Viability/Cytotoxicity Assay Kit (Biotium #30002), according to manufacturer’s guidelines. Cytoplasmic Calcein-AM identified live cells, while nuclear Ethidium Homodimer III (EthD-III) the dead cells. Cells treated with 0.1% w/v saponin for 10 min served as positive control for EthD-III staining. Experiments were performed in duplicates.

### Assessment of cell proliferation

The Click-iT EdU Cell Proliferation Kit for imaging (C10337 - ThermoFisher Scientific) was used to assess cell proliferation, according to the manufacturer’s instructions. Cells were incubated with EdU for 24 hrs in the presence of S protein (1 μg/mL - 5.8 nM) or PBS vehicle, and then analysed. Wherever required, cells were incubated with the anti-CD147 neutralising antibody (20 μg/mL). Experiments were performed in duplicates.

### 2D-Matrigel angiogenesis assay

Human CAECs were seeded on the top of Matrigel (Corning® Matrigel® Growth Factor Reduced Basement Membrane Matrix, cat# 356231) either in monoculture (4,000 cells/well) or in coculture with PCs (4,000 CAECs + 1,500 PC/well), using Angiogenesis μ-Slides (IBIDI, UK) and growth factors free medium. The S protein (1 μg/mL - 5.8 nM) or PBS vehicle were added to the system. Images were taken after 5 hrs using an inverted Leica microscope equipped with a 5x objective. The total tube length per imaging field was measured using ImageJ. To assess the interaction between PCs and CAECs, PCs were labelled with the red fluorescent tracker *Vybrant*™ *DiI Cell-Labeling Solution* (Invitrogen; dilution 1:1,000 in PBS, incubation for 5 min at 37°C followed by 15 min at 4°C). For experiments requiring the CD147 blockade, 100,000 PCs in a total volume of 100 μL were pre-incubated with the anti-CD147 antibody (20 μg/mL) for 1 hour at 20°C. Experiments were performed in triplicates.

### Wound closure migration assay

Cells were seeded into 96-well plates. A scratch was produced in confluent PCs in the centre of each well using a 20 μL tip. Cells were washed with PBS to remove detached cells and incubated with EBM2 medium under FBS and growth factors deprivation. Cell proliferation was inhibited using hydroxyurea (2 mM). Where required, cells were pre-incubated with the anti-CD147 antibody (20 μg/mL) for 1 h at 37°C. The S protein (1 μg/mL - 5.8 nM) or PBS vehicle were added to the system. Images were snapped at baseline and after 16 hrs, using an inverted Leica microscope equipped with a 5x objective. The wound area was measured, and the percentage of wound closure calculated. Experiments were performed in 4 to 5 replicates.

### Assessment of ERK1/2 phosphorylation

For detection of ERK1/2 phosphorylation in cardiac PCs, either western blotting or the ERK1/2 ELISA® Kit (Abcam ab176660) were employed. When required, PCs were pre-incubated with the anti-ACE2 (20 μg/mL, as described before(7)) or anti CD147 (20 μg/mL) antibodies for 1 hat 37 °C. Cells were exposed to the S protein (1 μg/mL - 5.8 nM) or PBS vehicle for 1 h at 37°C.

### CD147/Basigin (*BSG*) silencing in cardiac PCs

Opti-MEM media (ThermoFisher Scientific, UK) and Lipofectamine RNAiMAX (Invitrogen, UK) were used to transfect PCs with On-target plus *BSG* SMARTpool (a mixture of 4 siRNA in a single reagent, L-010737-00-0005, Dharmacon, Horizon Discovery, UK). The siGENOME Non-Targeting siRNA Pool # 1 (Dharmacon, Horizon Discovery, UK) was used as a control (final concentration 25nM for both). The transfection reagent was removed after 6 hours and replaced with fresh EGM2 medium. On day 3 and 4 post-transfection, cells were used for functional assays. RNA and total cell lysates were collected, and silencing was confirmed using qPCR (below), ICC and western blotting.

### Gene expression analysis by real-time qPCR

Extracted total RNA was reverse-transcribed into single-stranded cDNA using a High-Capacity RNA-to-cDNA Kit (ThermoFisher Scientific). The RT-PCR was performed using first-strand cDNA with TaqMan Fast Universal PCR Master Mix (ThermoFisher Scientific). TaqMan primer-probes were obtained from ThermoFisher Scientific (*BSG* Hs00936295_m1; housekeeping gene *UBC* Hs00824723_m1). Quantitative PCR was performed on a QuantStudio™ 5 System (ThermoFisher). All reactions were performed in a 10 μL volume in triplicate, using 7.5 ng cDNA per reaction. The mRNA expression levels were normalised against UBC and determined using the 2^−ΔΔCt^ method.(37)

### Collection of PC conditioned medium

Confluent PCs were maintained for 24 hrs in serum- and growth factors-free medium. When required, cells were pre-incubated with the anti-CD147 antibody for 1 h (20 μg/mL). The S protein (1 μg/mL - 5.8 nM) or PBS vehicle were added for 24 hrs. Conditioned media were collected and centrifuged for 10 min at 1,000 *g*, at 4°C, and stored at −80 °C until use.

### Cytokines and chemokines array

A panel of 105 cito/chemokines was assessed in the conditioned medium of cardiac PC treated with either the Spike protein or PBS vehicle, using the Human XL Cytokine Array kit (R&D, ARY022B). Each membrane was incubated with 750 μL conditioned medium. The assay was carried out according to manufactures instructions. Blot densitometry data for Spike-treated PCs are reported as fold-change versus vehicle.

### Measurement of cytokines using ELISA

Monocyte chemoattractant protein-1 (MCP1/CCL2, R&D DY279), tumour necrosis factor (TNFα, R&D DY210), interleukin-6 (IL-6, R&D DY206), interleukin-1 beta (IL-1β, R&D DLB50) were measured in PC conditioned medium using ELISA. Cell protein extracts were collected using RIPA buffer and the total amount of protein was quantified for normalisation of secreted factors (BCA assay). Secreted factors were expressed as pg secreted protein per 100 μg of total cellular protein.

### Effect of PC secretome on EC apoptosis

CAECs were seeded in 96-well plates. After 24 hrs, cells were incubated for additional 24 hrs with the PC conditioned medium diluted 1:2 with fresh EBM2 growth factors-free medium, with a final serum concentration of 1%. CAECs were also incubated with the S protein (1 μg/mL - 5.8 nM) or PBS vehicle for 24 hrs. Cell apoptosis was assessed using either the CaspaseGlo 3/7 assay (G8090, Promega, UK) according to the manufacturer’s instructions (N=4 replicates), or the Calcein-AM/EthD-III viability kit (in duplicates). EC death was measured either as caspase activity (relative luminescence units) or as % of EthD-III-positive cells, and finally expressed as fold changes versus the control vehicle group.

### Statistical analyses

Data were analysed using Prism version 8.0 and expressed as individual values and as means ± standard error of the mean (SEM). Statistical differences were determined using unpaired T-tests or 1-way or 2-way ANOVAs as appropriate. Non-parametric tests were applied. Statistical significance was assumed when *P* ≤ 0.05.

## RESULTS

### Cardiac PCs express SARS-CoV-2 S protein receptors

Immunohistochemistry analyses of human hearts showed that a subset of PDGFRβ^+^ PCs surrounding the heart microvasculature express the S protein receptors ACE2 and CD147 *in situ* (**Figure 1A-D**).

**Figure 1.**
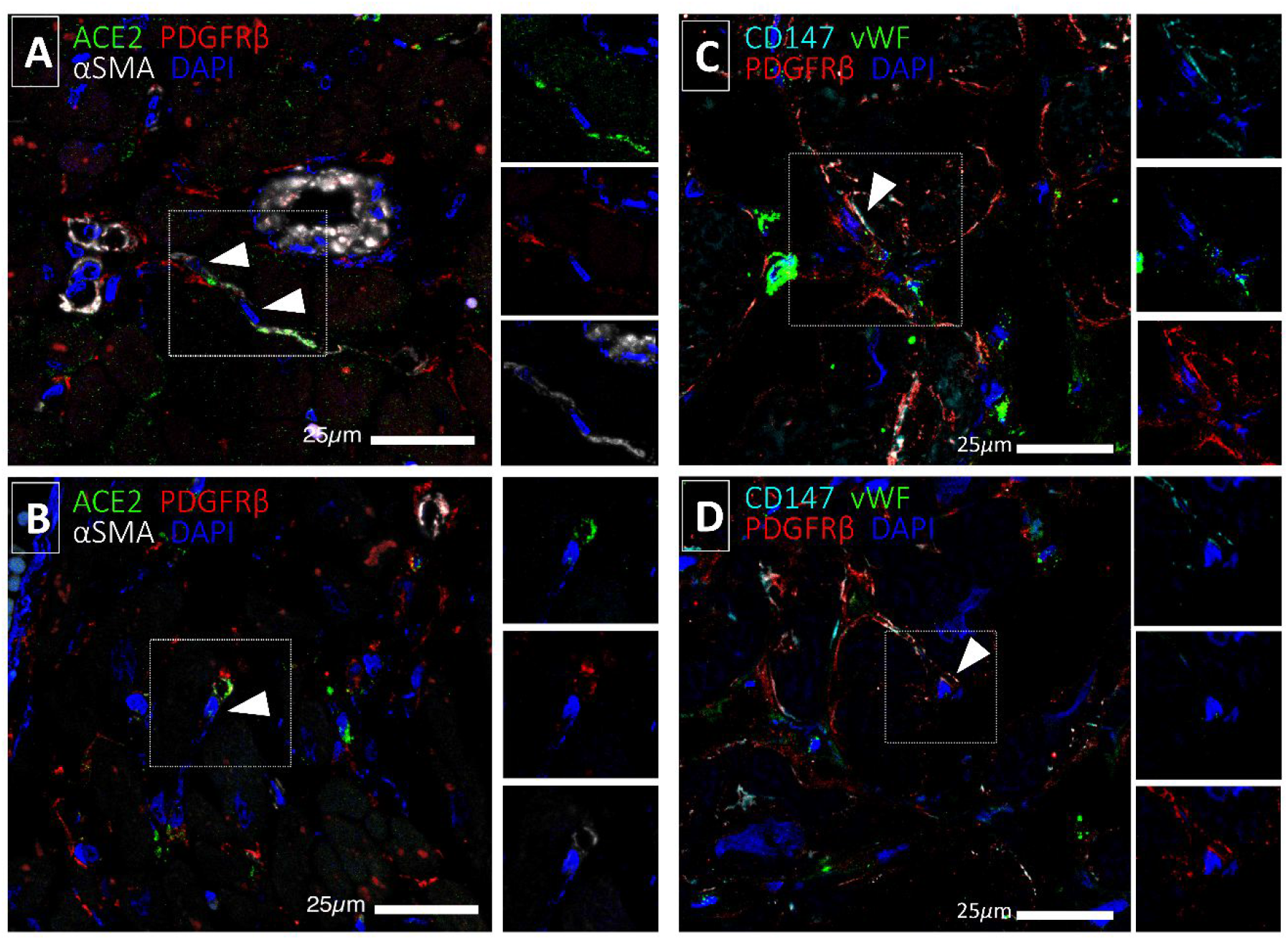
Expression of SARS-CoV-2 receptors in human cardiac PCs *in situ*. Immunofluorescence stainings of human hearts showing that a subset of Platelet Derived Growth Factor Receptor Beta (PDGFRβ)+ cardiac PCs express ACE2 **(A&B)** and CD147 **(C&D)**. Arrowheads point to PCs. Smooth muscle alpha-actin (αSMA) labels PCs and arterioles vascular smooth muscle cells (A&B), while von Willebrand Factor (vWF) recognises ECs (C&D).

We then confirmed the expression of S receptors in expanded cardiac PCs *in vitro*. Primary cultures of PCs were derived from myocardial leftovers of patients undergoing heart surgery, as previously reported.(22, 31) After expansion, PCs showed the characteristic spindle-shape and expressed the typical mural cell antigens NG2 and PDGFRβ, whilst being negative for the fibroblast marker PDGFRα (**Figure 2A**). Immunocytochemistry showed that PCs express the major SARS-CoV-2 receptor ACE2 and TMPRSS2, required for proteolytic activation of the S protein(7) (**Figure 2B**). Calu-3 and VeroE6/ACE2/TMPRSS2 cells were used as positive controls. Western blotting further indicated that cardiac PCs express considerably lower levels of both ACE2 and TMPRSS2 than control cells (**Figure 2C**). PCs also express CD147 (**Figure 1D&E**). For the last antigen, primary human coronary artery ECs (CAECs) were used as reference control.

**Figure 2.**
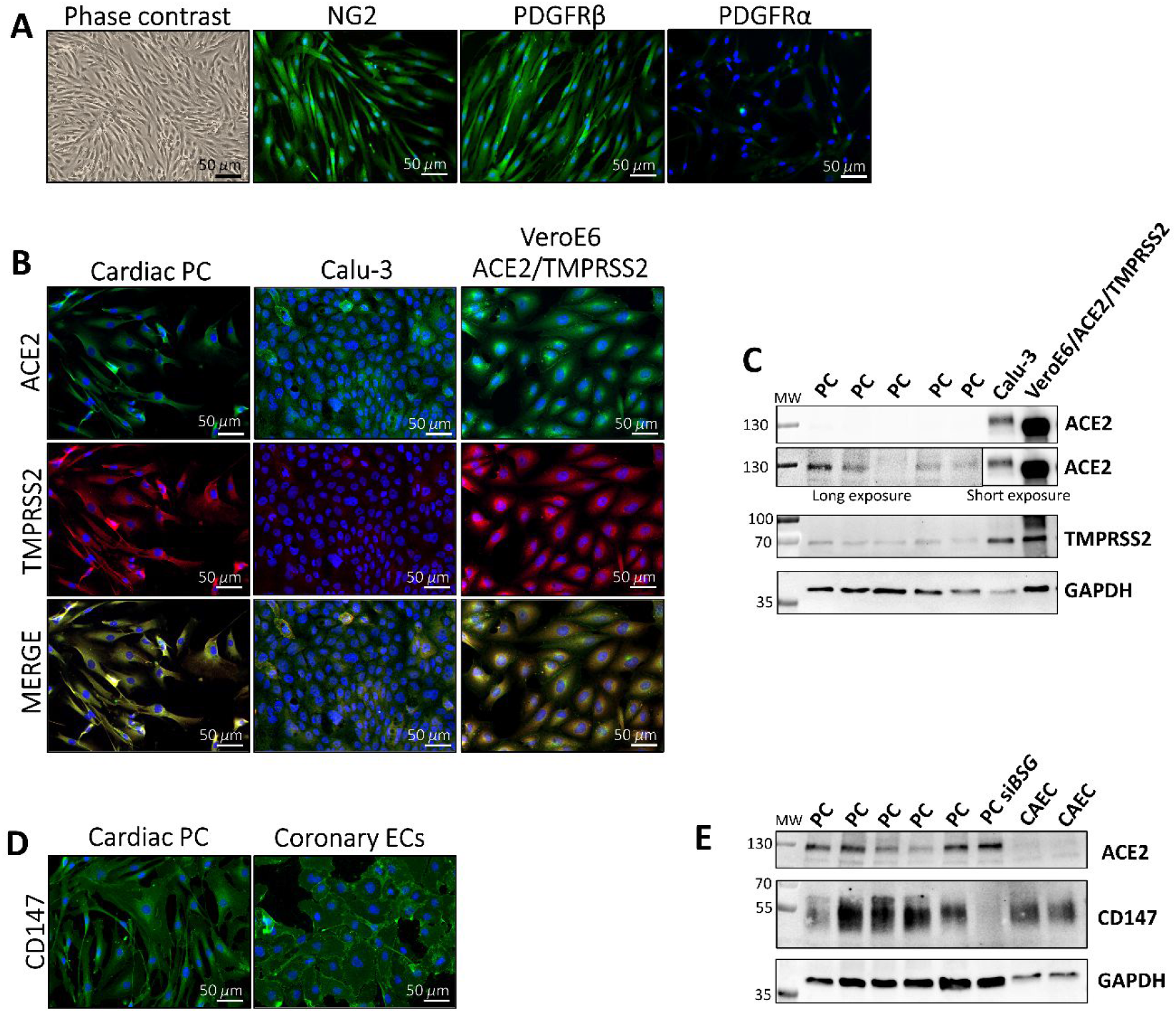
Expression of the SARS-CoV-2 receptors ACE2 and CD147, and activating protease TMPRSS2, in cultured cardiac PCs. **(A)** Immunofluorescence images show the characteristic antigenic phenotype. In green fluorescence, the antigens as indicated. In blue nuclei (DAPI). NG2: Neural/glial antigen 2. **(B)** Expression of ACE2 (green) and TMPRSS2 (red) in cardiac PCs and control cells assessed by immunostaining. In blue nuclei (DAPI). Calu-3: human lung epithelial cell line. *VeroE6/ACE2/TMPRSS2*: African green monkey kidney cell line engineered to overexpress the human ACE2 and TMPRSS2. **(C)** Expression of ACE2 and TMPRSS2 assessed using western blotting. N=5 patient PCs, N=1 for cell lines. **(D)** Expression of CD147 (green) in cardiac PCs and control human coronary artery endothelial cells (CAEC) assessed by immunostaining. In blue nuclei (DAPI). **(E)** Expression of CD147 and ACE2 determined using western blotting. N=5 patient PCs. N=2 CAEC. *siBSG* = *BSG* (CD147) silencing, used as a negative control.

### Cardiac PCs are not permissive to SARS-CoV-2 infection *in vitro*

Next, we investigated whether SARS-CoV-2 infects cardiac PCs *in vitro*. We employed a SARS-CoV-2 isolate from early in the pandemic (REMRQ0001) and the alpha variant (B.1.1.7) (MOI 10 for both). Permissive Caco-2-ACE2 cells were used as a positive control. Twenty-four hrs postinoculation, immunostaining for the N protein and dsRNA documented a lack of replicative infection in PCs (**Figure 3**). These findings were further confirmed by a longer infection time (five days, data not shown).

**Figure 3.**
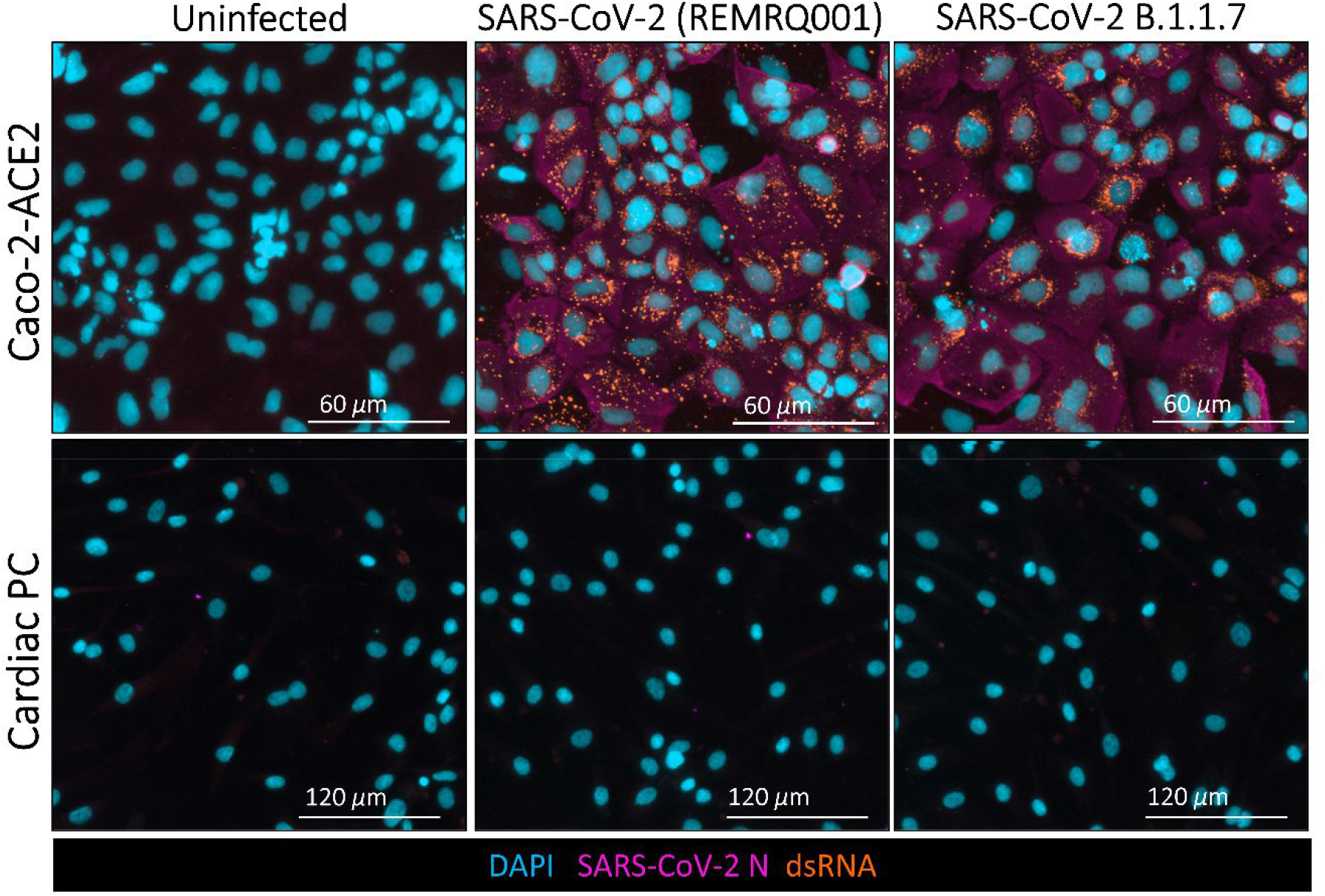
Cardiac PCs are not permissive to SARS-CoV-2 infection *in vitro*. Primary cardiac PCs (N=2 patients) and the Caco-2-ACE2 cell line were either mock-infected or inoculated with SARS-CoV-2 isolated early in the pandemic (REMRQ001) or the alpha variant (B.1.1.7) at a MOI 10 and incubated for 24 hours before immunostaining. Immunofluorescence images show SARS-CoV-2 nucleocapsid protein (N - magenta) and double-stranded RNA (dsRNA - orange) indicative of virus replication. Nuclei are stained with DAPI (blue).

### Presence of SARS-CoV-2 S protein in the peripheral blood of COVID-19 patients

We next explored the possibility that S protein cleaved from the virus by host cell proteases circulate in patients’ blood. Using a highly sensitive ELISA, we detected S protein in the serum of COVID-19 patients (33.5 ± 8.3 ng/mL) (**Figure 4A**). Although positive results were also recorded in the control group (12.7 ± 3 ng/mL), 18 out of 64 COVID-19 patients showed a S protein concentration higher than the 80th percentile of the control population (> 20 ng/mL) (**Figure 4A**). These events prevail in the patients group assessed between 5 - 10 days from the onset of COVID-19 symptoms (10/31 patients **- Figure 4B -** ANOVA *P*=0.056) and advanced age (60-75 years, 7/19 patients; > 75 years, 3/11 patients, **Figure 4C -** ANOVA *P*=0.047; 61-75 vs 20-45 years groups, *P*<0.05). The concentration of S protein in the different groups was uniformly distributed according to gender (data not shown).

**Figure 4.**
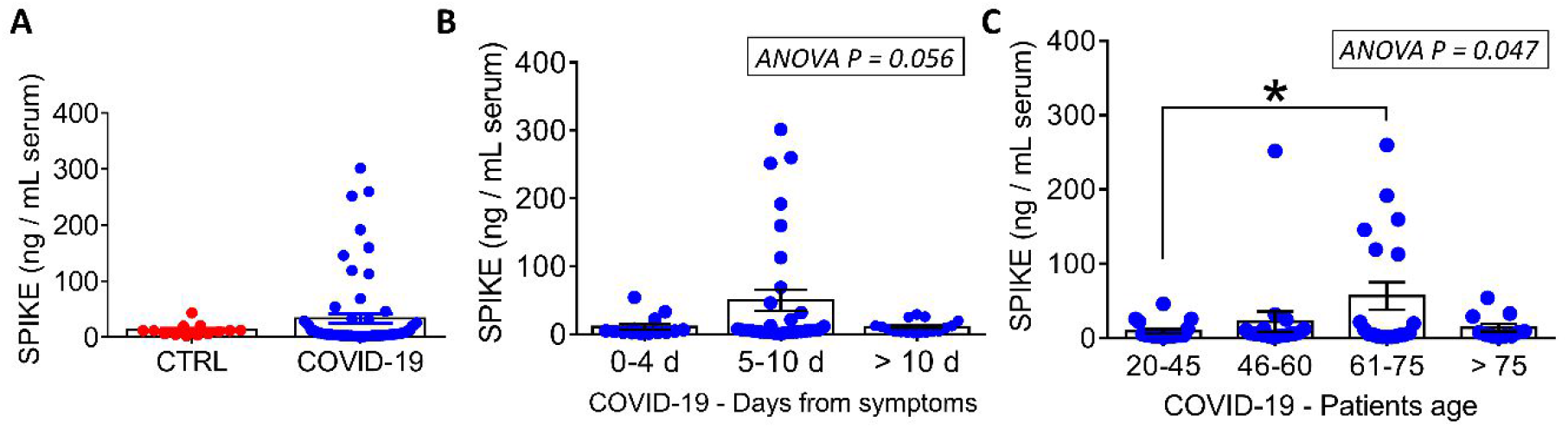
Measurement of S protein in the serum of COVID-19 patients and pre-pandemic controls. **(A)** Comparison of controls (N=14) vs COVID-19 patients (N=64). **(B)** Distribution of data according to the time from the onset of the disease symptoms. **(C)** Distribution of data according to the patients’ age. Graphs report individual values and means ± SEM. \**P*< 0.05.

### The SARS-CoV-2 S protein interacts with and causes dysfunction of cardiac PC

The viral S protein can engage with cellular receptors even in the absence of viral entry. Likewise, soluble S protein can potentially interact with host cell receptors. Therefore, we verified whether a recombinant extracellular domain of the S protein binds to cardiac PCs, thereby triggering intracellular molecular events and functional phenomena. The recombinant S protein was tagged with a hexahistidine (HIS) sequence for easy detection. Using Western blotting, we found bands corresponding to the HIS-tagged S protein in PCs exposed to the protein for an hour (**Figure 5A**). The recombinant S protein was used as a positive control.

**Figure 5.**
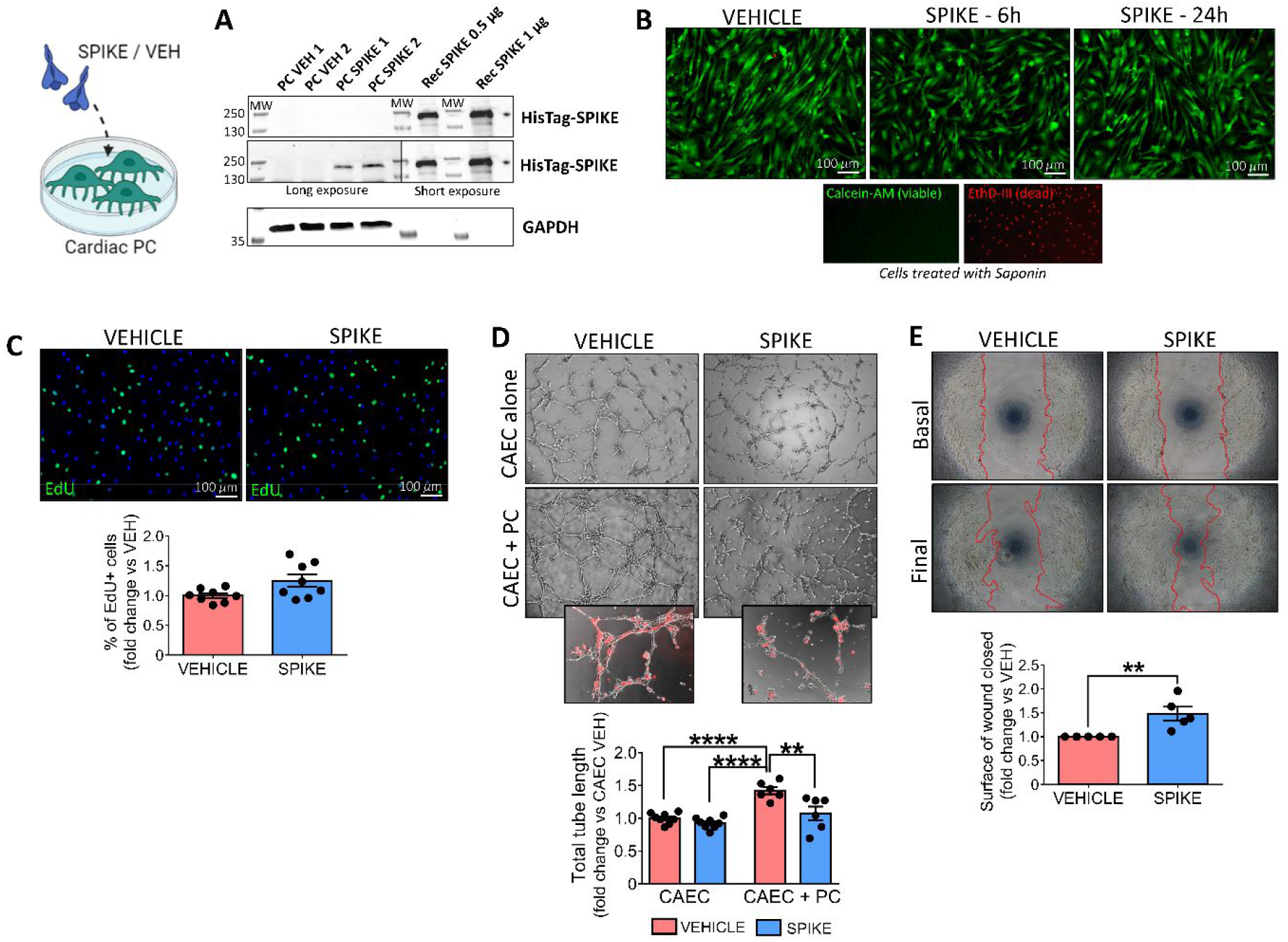
The SARS-CoV-2 recombinant S protein interacts with cardiac PCs and induces functional alterations. **(A) S protein interaction with PC receptors.** Western blotting analysis of PCs (N=2 patients) exposed to the S protein (SPIKE) or PBS vehicle (VEH) for 1 hour. The bands corresponding to the 6x-His-Tag recognise the His-tagged S protein. The purified S protein was used as a positive control. **(B) Cell viability.** Live cell imaging of PCs after 6 and 24 hrs incubation with the S protein. In green, Calcein-AM shows the cytoplasm of live cells. The red fluorescence of EthD-III (Ethidium Homodimer III) indicates the nuclei of dead cells (not detected). Saponin treatment, used as a positive control for dead cells, shows the nuclear staining of EthD-III in the absence of Calcein-AM. **(C) Cell proliferation**. PCs were exposed to the S protein or vehicle for 24 hrs, in the presence of EdU. Proliferation was measured as the % of EdU+ cells and data expressed as fold-change vs vehicle. Immunostaining shows EdU+ cells in green, nuclei (DAPI) in blue. N=8 patient PCs. **(D) Matrigel assay.** Coronary artery ECs (CAEC) and cocultures of CAEC + PCs were incubated on the top of Matrigel for 5 hrs, in the presence of the S protein or vehicle. PCs were labelled with the red fluorescent tracker CM-Dil to assess the interaction with ECs (small inserts). Graphs report the total tube length per imaging field, expressed as fold-change vs CAEC vehicle. N=6 patient PCs. **(E) Migration wound closure assay**. A scratch was created in confluent PCs and images taken at baseline. Cells were incubated with the S protein or vehicle for 16 hrs and final images were recorded. The surface of wound closure was calculated as % of the baseline area and expressed as fold-change vs vehicle. N=5 patient PCs. For all assays, the SARS-Cov-2 S protein was used at a concentration of 1 μg/mL. Graphs report individual values and means ± SEM. \*\**P* < 0.01, \*\*\*\**P* < 0.0001.

We then tested whether the S protein alters PC function. Results show that exposure of PCs to the S protein for 6 or 24 hrs did not affect either PC viability (**Figure 5B**) or proliferation (**Figure 5C**). In an *in vitro* angiogenic assay, the presence of PCs (identified by staining with a red fluorescent dye) increased the formation of CAEC networks (CAEC+PC vs CAEC monoculture, *P*<0.0001), with this response being diminished in the presence of the S protein (CAEC+PC Spike vs CAEC+PC vehicle, *P*<0.01). In contrast, the S protein did not inhibit network formation by CAECs in the absence of PCs (**Figure 5D**). Finally, in a wound closure assay, the S protein increased the motility of cardiac PCs (Spike vs vehicle, *P*<0.01) (**Figure 5E**).

### S protein-induced effects on cardiac PC function are CD147 dependent

The next step was to investigate the intracellular signalling triggered by the S protein. As shown in **Figure 6A**, PCs treated with the S protein had significantly increased levels of phospho-ERK1/2 (ratio P-ERK1/2 to total ERK1/2, Spike vs vehicle, *P*<0.05). A CD147 neutralising antibody (CD147AB) abolished this response (Spike vs Spike+CD147AB, *P*<0.05), while an antibody anti-ACE2 did not. As shown in **Figure 6B**, the CD147 blockade also prevented the S protein from inducing PC migration (Spike vs Spike+CD147AB, *P*<0.05) and inhibiting PC-CAEC network formation on Matrigel (Spike vs Spike+CD147AB, *P*<0.05, **Figure 6C**).

**Figure 6.**
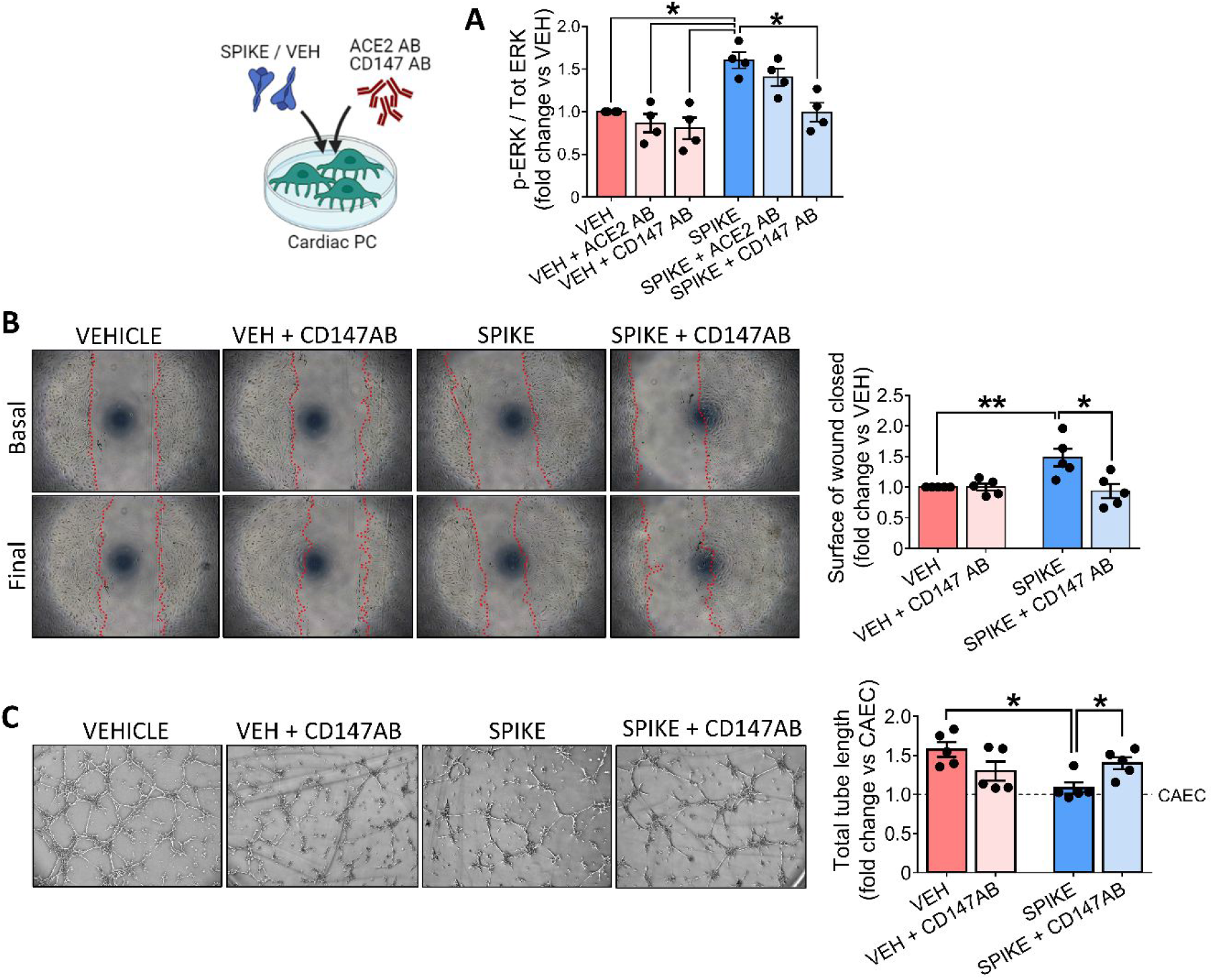
The SARS-CoV-2 S protein effects on cardiac PC function are CD147-dependent. **(A) Intracellular ERK1/2 phosphorylation/activation.** Cardiac PCs were cultured for 24 hrs under serum and growth factors deprivation and then exposed for 1 hr to the S protein (1 μg/mL) or vehicle. For receptor blockade, PCs were pre-incubated with antibodies anti-ACE2 or anti-CD147 for 1 hr in advance of the S treatment. The bar-graph reports the ratio between phospho-ERK1/2 (Thr202/Tyr204) and total ERK1/2, as measured by ELISA, and expressed as fold-change versus vehicle. N=4 patient PCs. **(B) Migration wound closure assay**. A scratch was created in confluent PCs and images taken at baseline. Where the blockade of CD147 was required, cells were pre-incubated with an antibody anti-CD147 for 1 hr. Then, the S protein (1 μg/mL) or vehicle were added to the system for 16 hrs. Final images were recorded. The surface of wound closure was calculated as % of the baseline area and expressed as fold-change vs vehicle. N=5 patient PCs. **(C) Matrigel assay.** Where the blockade of CD147 was required, cardiac PCs were pre-incubated with an antibody anti-CD147 for 1 hr. Afterwards, coronary artery ECs (CAECs) and cocultures of CAECs + PCs were incubated on the top of Matrigel for 5 hrs, in the presence of the S protein (1 μg/mL) or vehicle. Representative images of CAECs + PCs cocultures. The bar-graphs indicate the total tube length per imaging field, expressed as fold-change vs CAEC in single culture (dotted line at *y*=1). N=5 patient PCs. Graphs indicate individual values and means ± SEM. \**P*< 0.05, \*\**P*<0.01.

To confirm the leading role of CD147 in determining the PC response to the S protein, we repeated the functional assays using cardiac PCs that were silenced for CD147/Basigin (*BSG*). The silencing efficacy was confirmed using immunocytochemistry (**Figure 7A**), qPCR (**Figure 7B**), and western blotting (**Figure 7C**). We also verified that CD147 knockdown did not affect the cell viability (**Figure 7D**). When exposed to the S protein, *BSG*-silenced PCs showed less phosphorylation/activation of ERK1/2 than control cells (*P*<0.01, **Figure 7E**). Moreover, *BSG* silencing prevented the increase in PC motility in the presence of the S protein (si*BSG* Spike vs siCTRL Spike, *P*<0.05, **Figure 7F**) and rescued the pro-angiogenic activity of PCs on Matrigel (**Figure 7G**).

**Figure 7.**
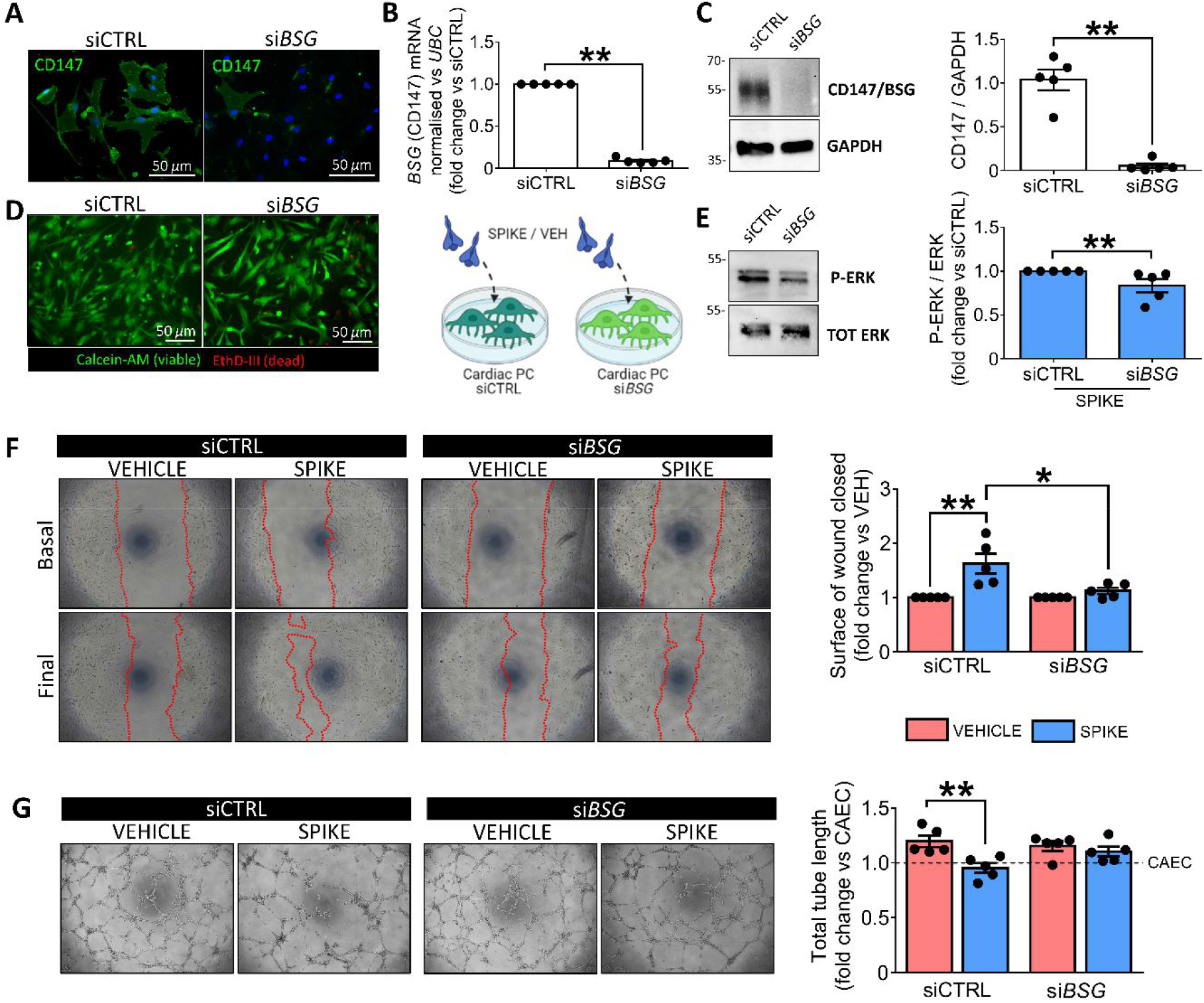
Basigin *BSG/CD147* silencing demonstrates that the SARS-CoV-2 S protein effects on cardiac PC function are CD147-dependent. **(A-C) *BSG* silencing in PCs.** CD147 protein knockdown in cardiac PCs was obtained using a pool of four small interfering RNA (siRNA). The silencing efficacy was confirmed using immunocytochemistry (A), qPCR (B) and western blotting (C). si*BSG*: *BSG* silencing. siCTRL: non targeting siRNA control. **(D) Cell viability post-silencing**. In green, Calcein-AM shows the cytoplasm of live cells. The red fluorescence of EthD-III (Ethidium Homodimer III) indicates the nuclei of dead cells. **(E) Intracellular ERK1/2 phosphorylation/activation in si*BSG* PCs**. Cardiac PCs were cultured for 24 hrs under serum and growth factors deprivation and then exposed for 1 hr to the S protein (1 μg/mL) or vehicle. The bar-graph reports the ratio between phospho-ERK1/2 and total ERK1/2, as measured by western blotting, expressed as fold-change vs siCTRL. N=5 patient PCs. **(F) Migration wound closure assay**. A scratch was created in confluent PCs and images taken at baseline. The S protein (1 μg/mL) or vehicle were added to the system for 16 hrs. Final images were recorded. The surface of wound closure was calculated as % of the baseline area and expressed as foldchange vs vehicle. N=5 patient PCs. **(G) Matrigel assay.** Coronary artery ECs (CAECs) and cocultures of CAECs + PCs were incubated on the top of Matrigel for 5 hrs, in the presence of the S protein (1 μg/mL) or vehicle. Representative images of CAECs + PCs cocultures. The bar-graph indicates the total tube length per imaging field, expressed as fold-change vs CAEC in single culture (dotted line at *y*=1). N=5 patient PCs. Graphs indicate individual values and means ± SEM. *\*P*< 0.05, \*\**P*<0.01.

### S protein-primed PCs secrete pro-inflammatory cytokines and induce EC death

Finally, we assessed whether the S protein triggers the production of a pro-inflammatory secretome in cardiac PCs, resembling the cytokine storm.(38) The analysis of PC conditioned media using a human cytokine/chemokine protein array revealed ten factors significantly regulated by the S protein treatment (Spike vs vehicle, *P*<0.05, **Fig 8A**). Among these, the potent pro-inflammatory cytokines TNFα, IL-6, and MCP1 were upregulated (average fold-change Spike vs vehicle ≥ 2, *P*<0.05, **Fig 8A**). We confirmed these findings using quantitative ELISAs, which also revealed the S protein induced PCs to secrete larger amounts of IL-1β (Spike vs vehicle, *P*<0.05, **Fig 8B**). The CD147 neutralisation failed in preventing all these alterations (Spike vs Spike+CD147AB, not significant - **Fig 8B**).

**Figure 8.**
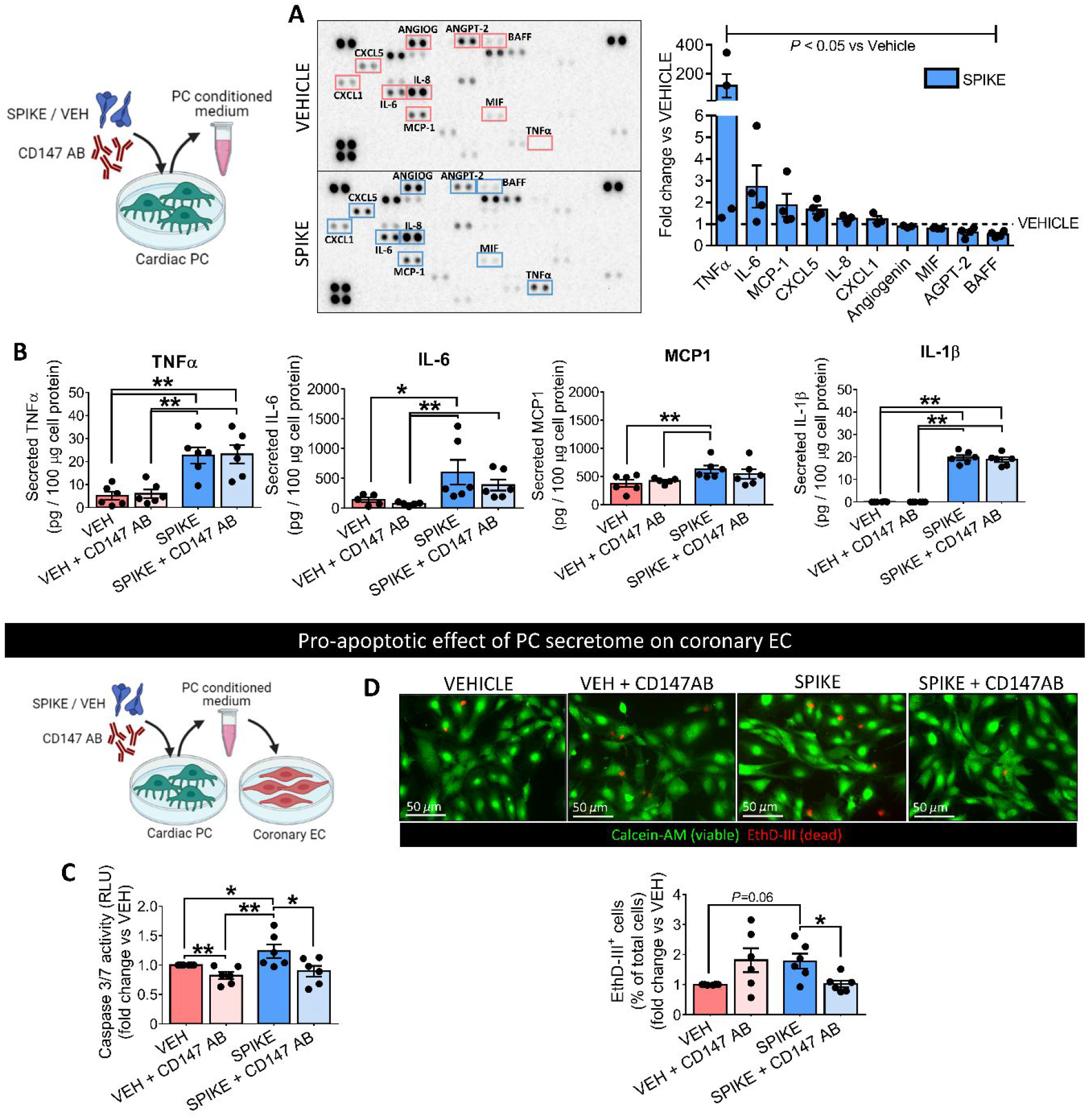
The secretome of S protein-challenged cardiac PCs is enriched in pro-inflammatory factors and induces EC death. **(A) Cytokine/chemokine protein arrays.** Cardiac PCs (N=4 patients) were incubated with the S Protein (1 μg/mL) or vehicle for 24 hrs. The conditioned medium was collected and analysed in the arrays. Per each factor, average pixel densities were measured and expressed as fold-change vs vehicle (dotted line at *y*=1). Ten factors were significantly regulated by Spike treatment (*P*<0.05 vs vehicle). **(B) Quantitative ELISAs.** Cardiac PCs (N=6 patients) were incubated with the S Protein (1 μg/mL) or vehicle for 24 hrs. For receptor blockade, PCs were preincubated with an anti-CD147 antibody for 1 hr in advance of the S treatment. The conditioned medium was collected and analysed using ELISA. Secreted factors were normalised against the total cellular proteins. **(C&D) The pro-apoptotic effect of cardiac PC secretome on coronary artery EC (CAEC) is prevented by CD147 blocking.** CAECs were incubated with the PC conditioned medium for 24 hrs, and cell death was evaluated by measuring Caspase 3/7 activity (**C**, relative luminescence units (RLU), early-stage apoptosis) and the % of cell nuclei positive for EthD-III (Calcein-AM/ Ethidium Homodimer III assay **- D -** positivity for EthD-III indicates irreversible cell death). Values are expressed as fold-change vs vehicle. N=6. Graphs show individual values and means ± SEM. * *P* < 0.05, ** *P* < 0.01.

Last, we checked whether the pro-inflammatory PC secretome harms EC viability. As shown in **Figure 8C**, exposure to the media from S protein-primed PCs induced the Caspase 3/7 activity in CAECs (Spike vs vehicle, *P* < 0.05), with this pro-apoptotic effect being reduced by the anti-CD147AB (Spike vs Spike+CD147AB, *P* < 0.05). These data were further confirmed by the fluorescent staining for EthD-III, indicating irreversible cell death (**Fig 8D**). Conversely, the S protein did not cause a direct pro-apoptotic effect on CAECs (data not shown).

## DISCUSSION

Our study provides novel *proof-of-concept* evidence for S protein to cause molecular and functional changes in human vascular PCs, both dependent and independent of the CD147 receptor (summarised in **Figure 9**).

**Figure 9.**
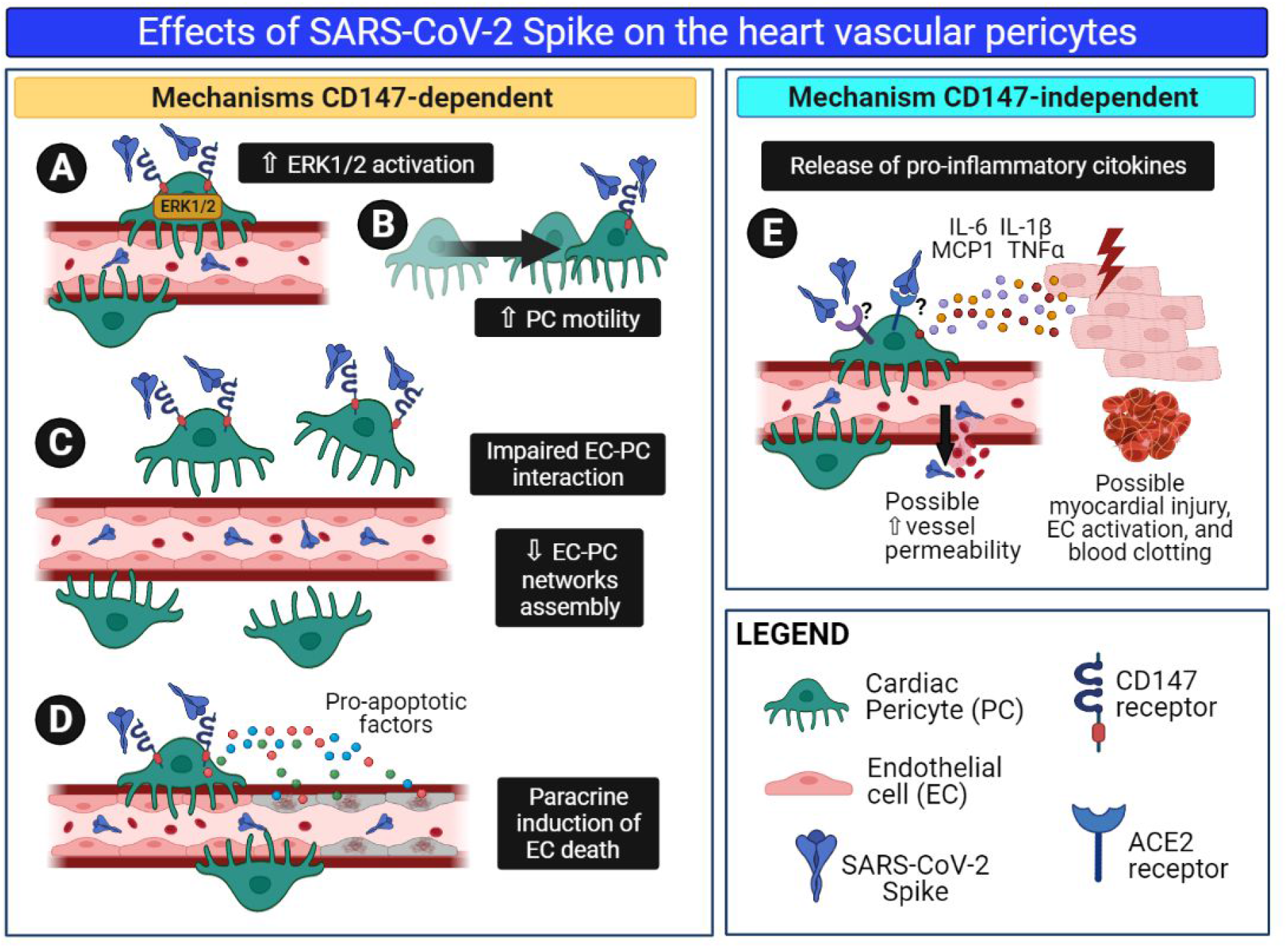
The SARS-CoV-2 S protein alters cardiac pericyte function. Schematic summary of the research. We hypothesize that in patients with acute COVID-19, S protein molecules are cleaved from the virus particle and released from the respiratory system into the bloodstream. Through the circulation, isolated S protein fragments reach all organs of the body, including the heart. Here, the interaction of the S protein with the CD147 receptor on cardiac PCs triggers the ERK1/2 signalling **(A)** and provokes PC dysfunction, including increased cell motility **(B)** and decreased angiogenic activity in cooperation with coronary ECs **(C)**. In addition, the S protein-CD147 interaction prompts cardiac PCs to release pro-apoptotic factors, which cause EC death **(D)**. Finally, through a mechanism CD147-independent, the S protein induces PCs to release pro-inflammatory cytokines, which include MCP-1, IL-6, IL-1β, and TNF-α **(E)**. These cytokines can damage neighbouring cardiomyocytes and activate ECs, potentially promoting blood to clot and increasing vascular permeability.

The SARS-CoV-2 virus particle consists of the three structural proteins: S, membrane (M) and envelope (E), embedded in a lipid bilayer surrounding a helical nucleocapsid comprised of the viral genomic RNA bound to the N phosphoprotein. While the M and E proteins are involved in viral assembly, the S protein mediates cell entry following priming/activation by host cell proteases, primarily TMPRSS2.(7) Moreover, the S protein activates the Raf/MEK/ERK signal transduction pathway in host cells.(6) Manipulation of the host ERK1/2 signalling pathway is reportedly instrumental to viral replication,(6, 39) and the induction of cyclooxygenase-2, a prostaglandin synthetase involved in inflammation.(40) Pharmacological inhibition or knockdown of ERK1/2 by small interfering RNAs suppressed coronavirus replication.(39) Therefore, drugs that block the binding of SARS-CoV-2 S protein to cell receptors and/or inhibit downstream signalling pathway may be potential candidates for the treatment of COVID-19.

Our study provides novel insights into the mechanism used by the virus to cause vascular damage. The classical route of infection starts with the multifunctional S protein binding to cell receptors, which opens the path to virus entry and the subsequent manipulation of the host intracellular machinery to cause cell dysfunction. Yet, here we show that cardiac vascular PCs are not infected by SARS-CoV-2, at least *in vitro*. This emerged using SARS-CoV-2 isolated early in the pandemic in the UK and the more transmissible SARS-CoV-2 alpha variant (B.1.1.7 lineage) that spread in the UK in early 2021. We hereby describe an alternative non-infectious mechanism triggered by the S protein alone in cardiac PCs. In support of this hypothesis, we found soluble S protein in the serum of COVID-19 patients. In particular, ~ 33% of patients in the advanced age group and during the acute phase of the disease presented elevated serological concentrations of S protein. It is possible that these molecules, cleaved from the virus particle by cellular proteases, spread from the respiratory system into the circulation. The presence of S protein in the peripheral blood only in the window between 5 and 10 days is compatible with subsequent neutralisation by blocking antibodies produced by immune cells. Although we cannot exclude that the antibody also detected S protein embedded in the whole virions, earlier studies showed that only low viral RNA copy numbers, below the threshold for viral infectivity, were detectable in a minority ( ~ 12%) of blood samples collected during acute infection.(41) Therefore, evidence suggests that shed S protein is far more abundant than whole SARS-CoV-2 particles in patients’ sera. Previously, another group identified S protein in patients’ blood.(42) In our assay, the non-specific positivity to S protein detected in pre-pandemic control sera could be explained by the sequence homology between some regions of the S protein and other human proteins, similar to what was previously described for the SARS-CoV-1.(43) To overcome this limitation, we only considered biologically relevant the concentrations of S protein higher than the 80^th^ percentile of the control group.

A few reports showed that the S protein alone, given to rodents either as a soluble molecule or presented with a carrier, exerts microvascular damage and induces hyperinflammation.(15, 44–46) Moreover, it was suggested a possible role for soluble S protein fragments in triggering blood clotting.(47, 48) These findings corroborate an essential biological role for the S protein beyond the presence of the whole viral particles. In addition, soluble S protein may remain engaged with cellular receptors for longer than the whole coronavirus, resulting in prolonged stimulation of intracellular signalling. Our *in vitro* studies collected several findings indicating that the S protein modifies PC biological properties. The concentration of S protein employed for the experiments is similar to those of other ligands known to activate ERK1/2 in PCs (EGF - 0.83 nM, and bFGF - 0.6 nM, former data from our group).(31) S protein-induced activation of PC migration, and inhibition of interactive cooperation between PCs and CAECs in angiogenesis assays, are both suggestive of a dysfunctional PC phenotype. In fact, the detachment of PCs from the perivascular compartment results in vulnerable capillaries prone to instability, pathological angiogenesis, and, ultimately, rarefaction. On the other hand, the S protein did not impinge upon PC viability. SARS-CoV-2 cannot replicate without the machinery of a host cell. Therefore, it could be counterproductive for the virus to kill the cell at the initial stage of the S protein engagement with host receptors. Interestingly, our experiments showed that the S protein does not alter CAEC viability and angiogenic capacity, thus suggesting a specific action of the S protein on PCs. The S protein also activated or enhanced the production of pro-inflammatory cytokines by cardiac PCs. MCP1, IL-6, IL-1β and TNFα are typical components of the cytokine storm associated with respiratory failure and high mortality in COVID-19 patients.(38, 49) This pro-inflammatory secretome could produce harmful paracrine effects on the surrounding vascular cells, as our experiment on CAEC apoptosis suggests. This mechanism can propagate functional alterations even to those cell populations which may not be directly infected by the virus, ultimately contributing to vascular disruption.

A recent report showed that the whole S1 subunit causes the phosphorylation/activation of MEK in human pulmonary vascular cells.(9) However, using only the ACE2 RBD failed to do so. Therefore, it was not clear if the signalling started from the ACE2 receptor.(9) The authors suggested that an alternative receptor, different from ACE2, might mediate the signalling of the S protein in vascular mural cells. Basigin/CD147, a plasma membrane protein associated with oligomannosidic glycans, has emerged as a novel receptor for SARS-CoV-2.(12) Supporting evidence from studies in epithelial and immune cells(50, 51) has been contradicted by investigations on CD147-transfected HEK 293 cells and biochemical experiments.(52, 53) These discrepancies may be reconciled with the requirement of coreceptors or post-translational modifications for the effective binding of the S protein to CD147. But supportive proof of the involvement of CD147 came from research *in vivo*. In a preclinical study with transgenic mice expressing human CD147, the administration of a neutralising antibody against the receptor successfully treated exudative pneumonia caused by SARS-CoV-2 infection.(51) And more importantly, an open-label clinical trial of meplazumab, a humanised therapeutic monoclonal antibody against CD147, showed striking improvements in COVID-19 patients.(54)

Our research confirms the data from single-cell sequencing studies showing cardiac PCs express the ACE2 mRNA transcript (data not shown).(19, 20) However, both ACE2 and TMPRSS2 protein levels in PCs were considerably lower than those of more permissive Calu-3 and VeroE6/ACE2/TMPRSS2 cells. This difference may account for the lack of virus infection in PCs. This observation is in line with recent reports showing that, *in vitro*, low ACE2 expression levels in myocardial stromal cells resulted in low susceptibility to viral infection,(55) whilst mesenchymal stromal cells were not infected due to the lack of ACE2/TMPRSS2.(56) Nonetheless, our data demonstrate that the CD147 receptor, and not ACE2, leads the S signalling in PCs. Indeed, CD147 blockade using a neutralising antibody or gene silencing, restrained the S protein from inducing ERK1/2 phosphorylation and rescued several functional features of PCs that were compromised by the S protein, including PC stability and pro-angiogenic activity. Finally, CD147 blockade protected CAECs from the paracrine apoptotic action of S protein-primed PCs. However, it failed to prevent the induction of multiple pro-inflammatory cytokines, thus suggesting the latter phenomenon involves mechanisms unrelated to the CD147 receptor in PCs.

In conclusion, although more investigation being needed to definitively prove the harmful effects of the S protein on the heart PCs and associated microvasculature *in vivo*, this work suggests that fragments of the S protein may elicit vascular cell dysfunction through CD147, independently from the infection. This mechanism has the potential to spread cellular and organ injury beyond the infection sites and may have important clinical implications. For instance, in patients with disrupted endothelial barrier and increased vascular permeability due to underlying diseases, such as hypertension, diabetes, and severe obesity, S protein molecules could easily spread to the PC compartment and cause, or exacerbate, microvascular injury. Blocking the CD147 receptor may help protect the vasculature of the most vulnerable patients from infection and the collateral damage caused by the S protein.

## Acknowledgements

We wish to acknowledge the members of the University of Bristol COVID19 Emergency Research Group (UNCOVER) for the continuous scientific support.

## Data availability

The data underlying this article will be shared on reasonable request to the corresponding authors.

